# Deep learning identifies heterogeneous subpopulations in breast cancer cell lines

**DOI:** 10.1101/2024.07.02.601576

**Authors:** Tyler A. Jost, Andrea L. Gardner, Daylin Morgan, Amy Brock

## Abstract

**Motivation:** Cells exhibit a wide array of morphological features, enabling computer vision methods to identify and track relevant parameters. Morphological analysis has long been implemented to identify specific cell types and cell responses. Here we asked whether morphological features might also be used to classify transcriptomic subpopulations within *in vitro* cancer cell lines. Identifying cell subpopulations furthers our understanding of morphology as a reflection of underlying cell phenotype and could enable a better understanding of how subsets of cells compete and cooperate in disease progression and treatment.

**Results:** We demonstrate that cell morphology can reflect underlying transcriptomic differences *in vitro* using convolutional neural networks. First, we find that changes induced by chemotherapy treatment are highly identifiable in a breast cancer cell line. We then show that the intra cell line subpopulations that comprise breast cancer cell lines under standard growth conditions are also identifiable using cell morphology. We find that cell morphology is influenced by neighborhood effects beyond the cell boundary, and that including image information surrounding the cell can improve model discrimination ability.

## Introduction

Tumors are comprised of cells with varying levels of genetic and non-genetic differences^1–7^. Viewing cancer as both an ecological and evolutionary^8,9^ process has led to the development of “adaptive therapies”^10,11^, which seek to contain the tumor rather than fully eliminate it. This perspective incorporates the inherent heterogeneity within cancer, as it seeks to address the subpopulations which proliferate when chemotherapy is applied. Heterogeneity, even within *in vitro* cell lines, has been well documented through single-cell sequencing methods^12–15^. These high-dimensional measurements are generally destructive, endpoint readouts and therefore can only provide snapshots in time, documenting the current state of the population. To better understand how relevant subpopulations react in response to various treatments and schedules, non-destructive methods that capture population dynamics in live cell samples are necessary.

Recently, image analysis using machine learning has demonstrated that cell-state properties such as metastatic invasion^16–18^, the induction of an epithelial to mesenchymal transition^19^, or the introduction of genetic perturbations^20^ can be detected through cellular morphology. Notably, deep learning has been effective at learning relevant biological features^21–24^. Many of these results have been demonstrated only using basic imaging techniques, such as phase contrast or brightfield microscopy. To further explore the capabilities of this approach, we asked whether more subtle cell-state differences are identifiable even within cell lines under standard culture conditions. Here we investigate whether cells which exhibit distinct RNA expression patterns can be discriminated based on cell morphology alone.

To explore the extent to which transcriptomic differences are expressed through cell morphology, we tested the ability of a deep convolutional neural network (CNN) to discriminate cancer cell heterogeneity in three model conditions (Figure 1a). In the first condition, we used heavily bottlenecked cell populations which had experienced high levels of doxorubicin exposure and compared them to healthy cells. In the second, we isolated and fluorescently labeled two transcriptomically-distinct subpopulations within the MDA-MB-231 cell line. In the third, we extended this to transcriptomic subpopulations in the MDA-MB-436 cell line. After identification and isolation of these populations, we developed an image segmentation pipeline that allowed us to perform instance segmentation on patches of images (Figure 1b). A CNN was then trained to discriminate between the two populations in each condition using only phase contrast imaging (Figure 1c). We demonstrate that cell morphology is not only reflective of even subtle differences in transcriptomic expression, but that this morphology is heavily influenced by cell-to-cell orientation and interaction.

**Figure 1.**
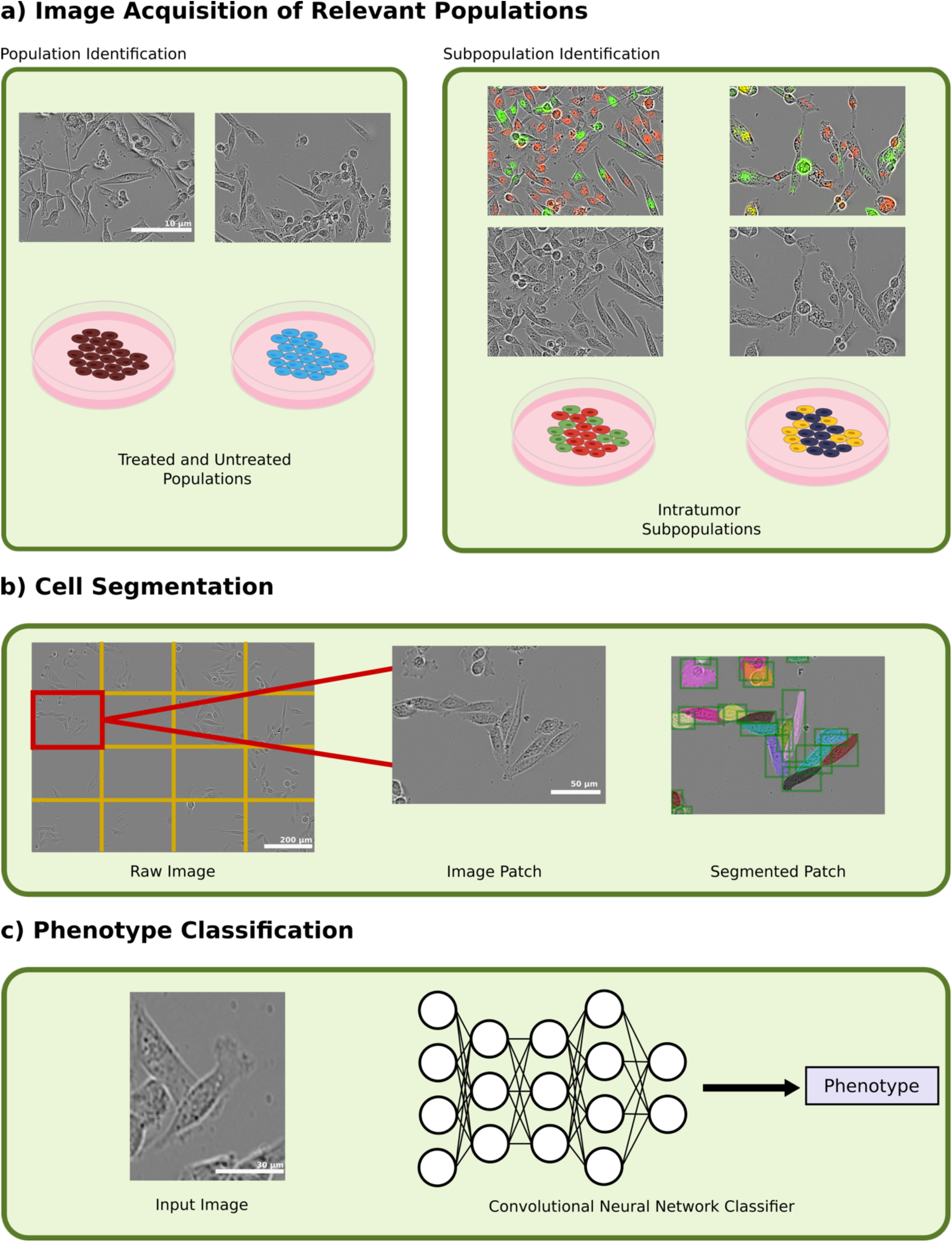
(a) Phenotype identification was done by first imaging treated and untreated populations of MDA-MB-231 cells and cocultured subpopulations in MDA-MB-231 and MDA-MB-436 cell lines at 20X magnification using phase contrast and fluorescent imaging. Images are contrast-enhanced for visualization purposes only. (b) Cell segmentation was then performed on each image by splitting the image into 16 patches and then segmenting using a Mask R-CNN segmentation network. (c) Input images were fed into a convolutional neural network classifier to that predicted each cell’s phenotype.

## Results

### Chemotherapy-treated Cells are Highly Identifiable

To establish a baseline for what is identifiable through cell morphology using our methodology, we explored how chemotherapeutic dosing of MDA-MB-231 cells affected their morphology before and after treatment. 50,000 MDA-MB-231 cells were seeded and treated with 550 nM of doxorubicin after 24 hours. After 48 hours, media containing the drug was removed and replaced with growth media. Cells were maintained in standard culture to reach a population size of 6 X 10^6^ cells/sample. We performed single-cell RNA sequencing (scRNA-seq) on both the treated population as well as an untreated control population. The full dataset was composed of 7502 treated and 3258 untreated cells after post-processing (Methods). Comparing overall RNA expression patterns revealed differences after treatment (Figure 2a), with the lowest observed level of correlation in gene expression between each condition (PCC −0.03, Supplementary Figure 1).

**Figure 2.**
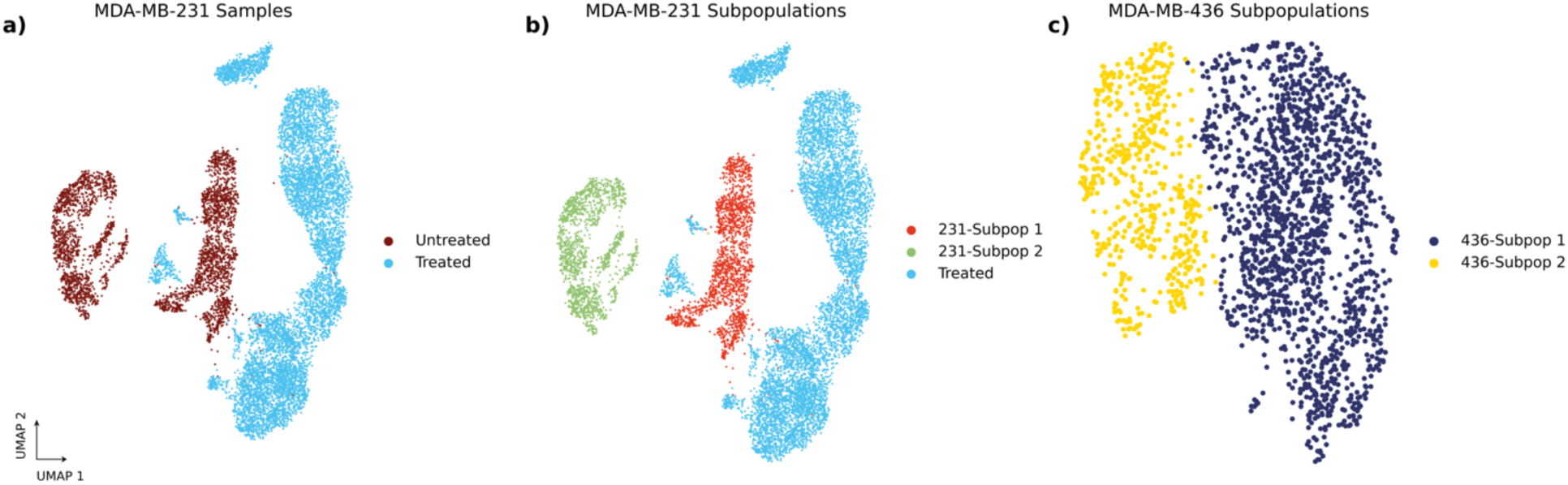
UMAP representations of (a) MDA-MB-231 Samples, (b) MDA-MB-231 subpopulations, and (c) MDA-MB-436 subpopulations. Subpopulations were identified as previously described previously^30^.

Next, we separately plated 60 wells of doxorubicin treated and 60 wells of control cells in 96 well plates and imaged at 20X magnification every 4 hours by phase contrast using Incucyte S3 (Sartorius) until cultures reached maximum confluency. We collected 77,325 instances of treated and 120,536 instances of untreated cells. Each image was then segmented using a custom-trained Mask R-CNN segmentation network (Methods). To determine cell identity, we used a pretrained Resnet-152 CNN^25^. Using transfer learning^26^, we were able to identify cells according to treatment status with an AUC of 0.95 (Figure 3). This is in line with current knowledge as it has been previously shown that doxorubicin induces an epithelial to mesenchymal^27,28^ transition and that this transition causes a distinct morphological shift.

**Figure 3.**
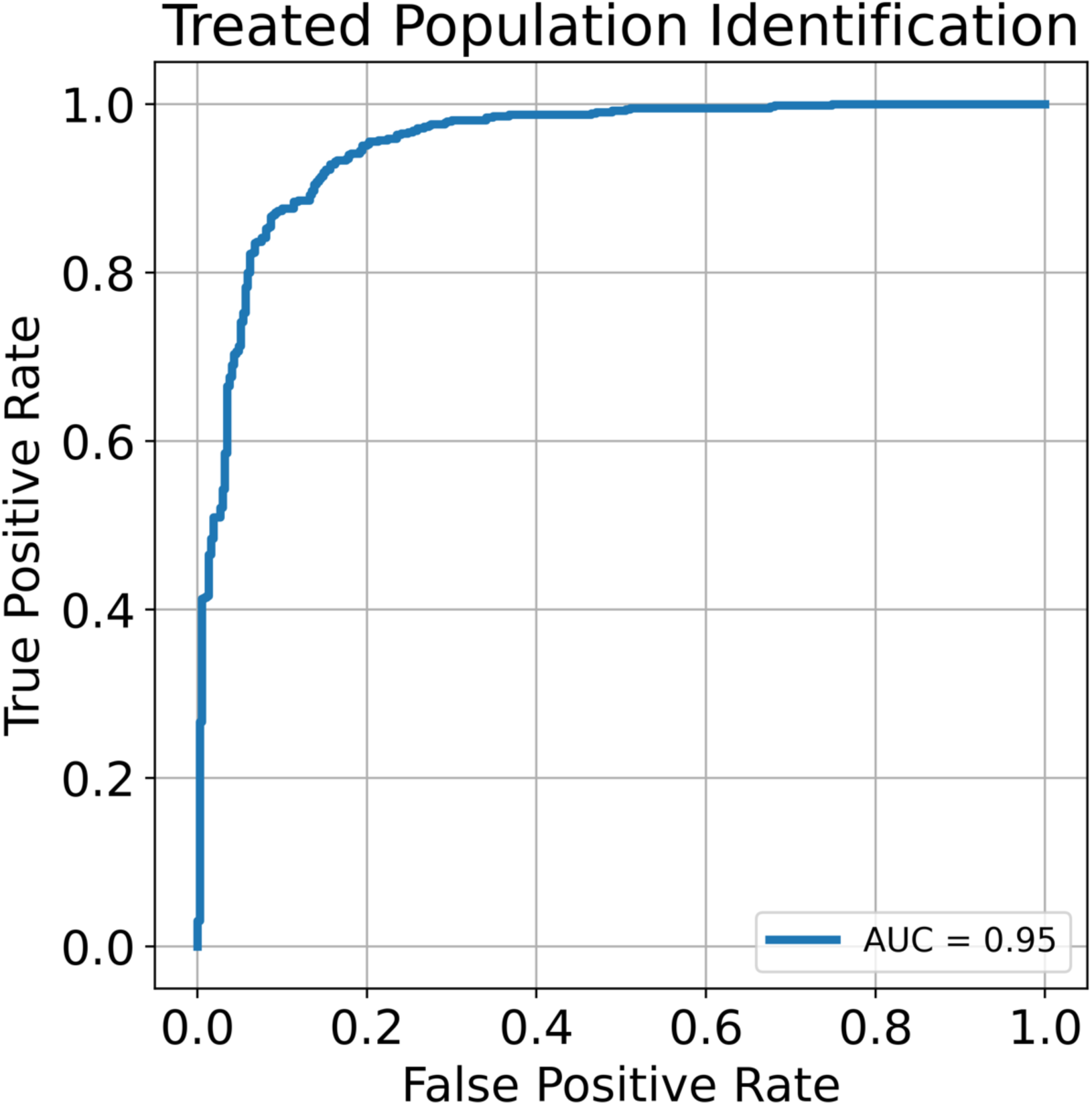
Receiver operating characteristics (ROC) curve for the untreated and treated populations.

### Identification of Transcriptomic Subpopulations in MDA-MB-231 cells

To determine whether it is possible to discriminate subsets of cells within a cell line, we examined scRNA-seq data of untreated MDA-MB-231 cells under standard growth conditions. We found that the MDA-MB-231 cell line consists of two transcriptomically-distinct populations (231-Subpop 1 and 231-Subpop 2) as identified by Leiden clustering^29^ (Figure 2b). Relative to the untreated and treated sample comparison, the two intra-cell line populations have more similar levels of gene expression (PCC=0.79, Supplemental Figure 1). We therefore asked whether these two subpopulations could be distinguished by the same approach of phase contrast imaging, segmentation, and phenotype classification.

In a previous study, we identified ESAM as a surface marker that differentiates the identified clusters^30^, with low expression in 231-Subpop 1 and high expression 231-Subpop 2. To establish ground truth knowledge of subpopulation identity during coculture imaging experiments, ESAM-separated subpopulations of MDA-MB-231 cells were established from parental cells with stable fluorescent nuclear labels on NLS-mCherry (ESAM-low, 231-Subpop 1) and NLS-GFP (ESAM-high, 231-Subpop 2) (Methods).

A 50/50 coculture of 231-Subpop 1 and 231-Subpop 2 cells was plated in 60 wells of a 96-well plate. They were imaged at 20X magnification every 4 hours with phase contrast and fluorescence imaging using an Incucyte S3. Because each population is fluorescently labeled based on its transcriptomic identity, we were able to label each cell’s phenotype after segmentation. In total we gathered 30,418 instances of cocultured 231-Subpop 1 and 63,758 instances of 231-Subpop 2 cells.

As before, we applied a Resnet-152 CNN to identify subpopulation identity, achieving an AUC of 0.74. We hypothesized that including neighborhood information such as cell-cell interaction could improve the ability of the network to correctly classify cells as belonging to 231-Subpop 1 or 231-Subpop 2. This environmental interaction is an under-utilized aspect of cell morphology that is not always applied when attempting to classify cells using only morphology. While there are large bodies of literature which have researched how natural density-dependent phenomena such as the Allee effect^31,32^ and contact inhibition^33–35^ influence tumor growth, cell morphology is often focused on elements such as texture and shape, as opposed to orientation and interaction with neighboring cells. To test this hypothesis, we designed several *in-silico* experiments to investigate the effect of neighborhood properties. In the first, we incrementally increased the bounding box around the input image to include neighboring cells. We tested several bounding box increases between 0 and 65 pixels (Figure 4a) and found that there exists an optimal range between pixel increases of 25 and 45 pixels which includes enough information about the orientation and interaction between cells (Figure 4b), but which does not obfuscate which cell is being identified. This resulted in a maximum AUC of 0.8.

**Figure 4.**
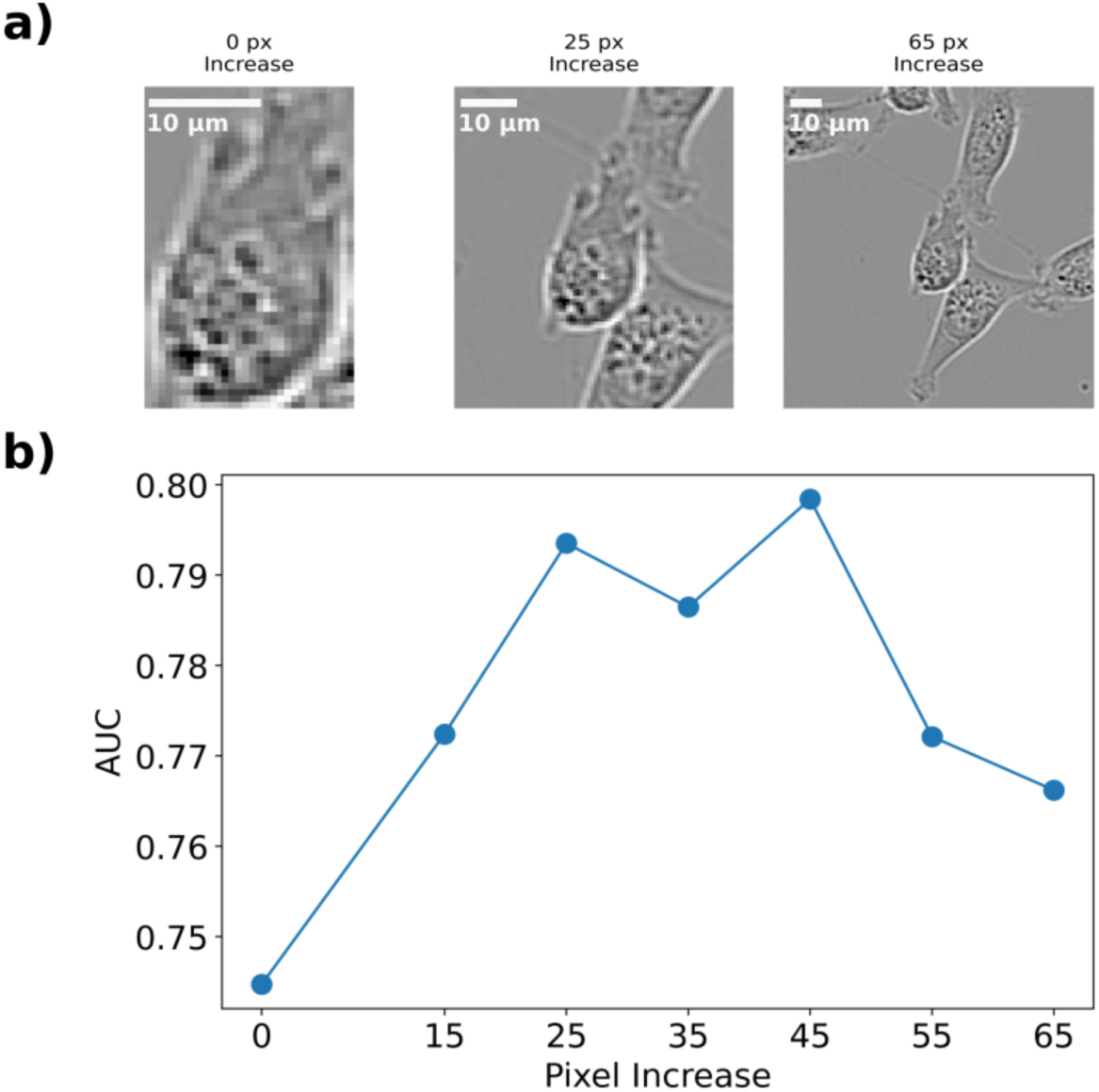
(a) Representative images of MDA-MB-231 cells with increasing bounding boxes. The segmented cells are centered and the bounding box around them is increased uniformly. (b) The AUC of the trained CNN versus the pixel increase around the cell. Increasing the bounding box increases the AUC until it reaches 0.80 around 25 pixels, at which point it plateaus then decreases.

To further test this hypothesis, we also designed *in-silico* experiments to selectively remove information (Figure 5a) using a 25 pixel bounding box increase. In the first, we completely blacked out the area around the cells. In the second, we whited out the cell itself. Intuitively, these experiments test the ability of the CNN to classify cell phenotype without information about background or without the information on texture. We find that the network classification ability diminishes with these limitations (Figure 5b), giving support to our hypothesis that including the environment around the cell is a contributing factor to phenotype identification in cell morphology.

**Figure 5.**
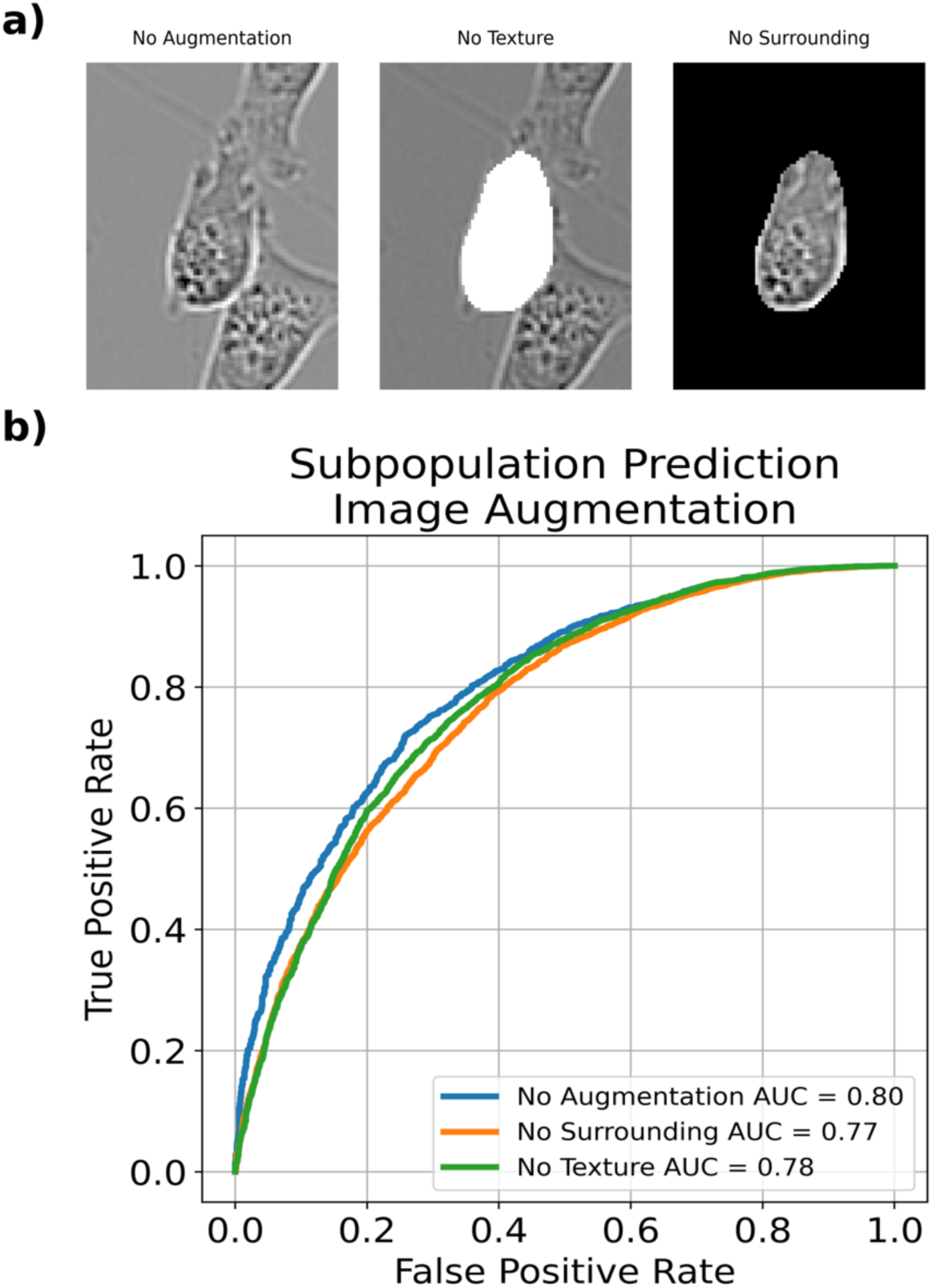
(a) Representative images MDA-MB-231 cells at a 25 pixel bounding box increased with no augmentation, a whited out cell for no texture, or a blacked out surrounding. (b) ROC plot for each given augmentation. Augmenting images revealed modest decreases in classification ability.

### Identification of Transcriptomic Subpopulations in MDA-MB-436 cells

After demonstrating that highly perturbed MDA-MB-231 cells as well as unperturbed subpopulations within the parental MDA-MB-231 population could be identified using deep learning, we questioned whether this phenomenon would extend to other cell models. Using unsupervised Leiden clustering, we identified the MDA-MB-436 cell line as another example of an *in vitro* breast cancer model consisting of two subpopulations with unique transcriptional patterns^30^. These transcriptomic subpopulations had the highest observed level of similarity (PCC=0.96, Supplemental Figure 1). Similar to the gene ESAM in the MDA-MB-231 cells, BST2 is a marker that differentiates each population. MDA-MB-436 subpopulations were sorted directly from a parent population using tetherin, the protein product of BST2. We grew isolated replicates for each subpopulation, then dyed each individually using a red cytoplasmic membrane dye for BST2 high cells (436-Subpop 1) and a green cytoplasmic dye for BST2 low cells (436-Subpop 2) (Methods).

We plated each isolated subpopulation in a 50/50 coculture in 60 wells within a 96 well plate. Each well was imaged at 20X magnification every 4 hours. We collected 86,924 instances of 436-Subpop 1 and 80,834 instances of 436-Subpop 2 cells. As before, cell phenotype was identified using the fluorescence of the segmented cell. Bounding box optimization was performed as above, with the max AUC= 0.75 at a 65 pixel increase. We found that the discrimination ability of the network increased as the bounding box increased, then dropped off similar to the MDA-MB-231 subpopulation identification (Figure 6).

**Figure 6.**
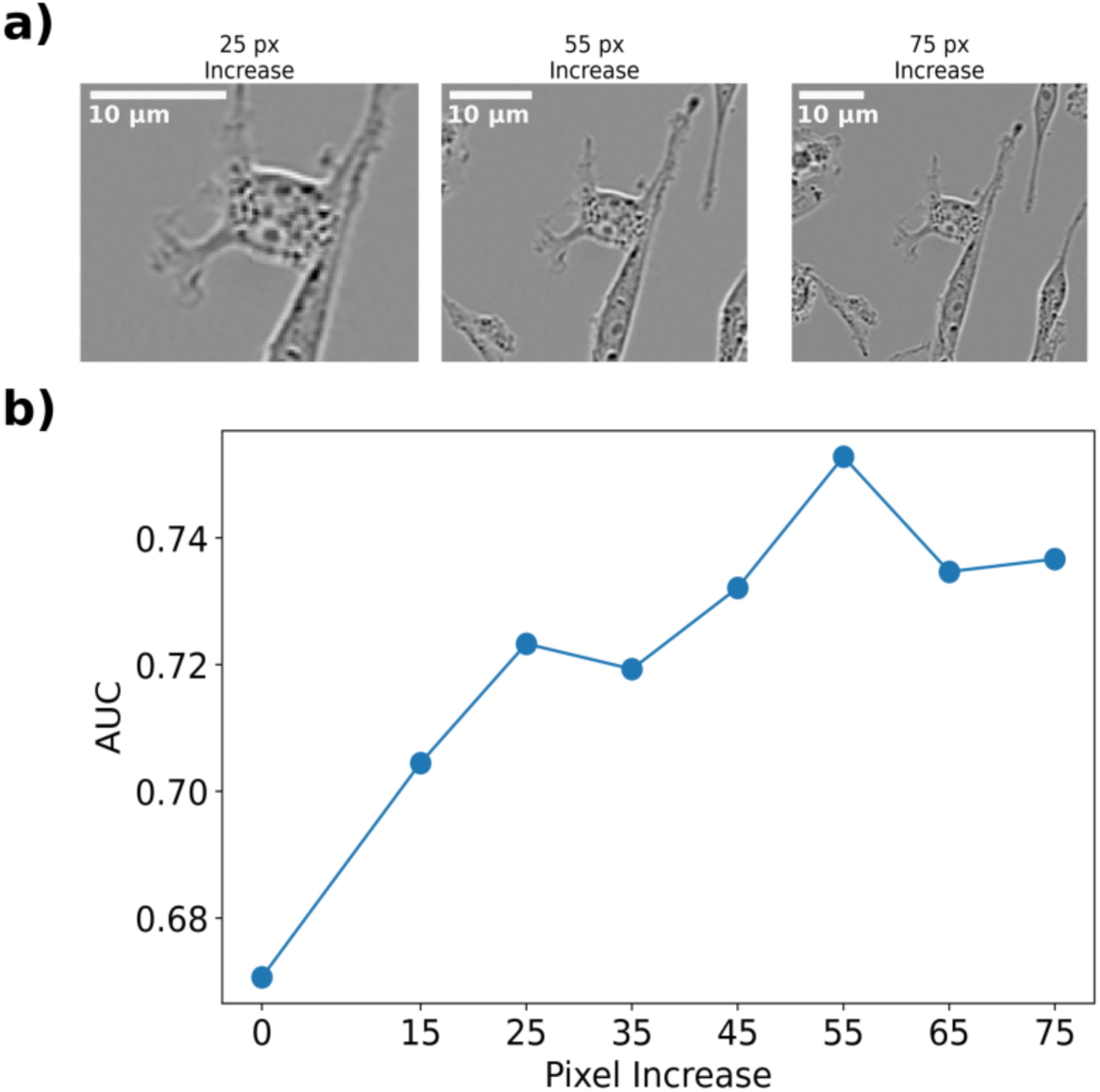
(a) Representative images of MDA-MB-436 cells with pixel bounding box increases. (b) The AUC of the trained CNN versus the pixel increase around the cell. As with the MDA-MB-231 cell line, the discrimination ability of the model increases with more neighborhood information, this time peaking with an AUC of 0.75 at a 55 pixel increase.

## Discussion

Cell morphology has long been used to identify properties of cells such as cell type and metastatic viability^18,36–40^. The heterogeneity that can exist within cell morphologies, however, presents significant challenges in correctly segmenting cells and identifying properties of interest. Deep learning has proved to be an ideal approach for classifying cells based on morphology^24^, as it has the ability to absorb a large corpus of data and learn relevant features.

In this study, we predicted cell phenotype using cell morphology from phase contrast images. We demonstrated that cell morphology reflects underlying transcriptomic differences and can be used to identify multiple types of in vitro populations. Additionally, we observed that including neighborhood context about cell orientation and interaction with other cells increased the accuracy of the model.

We tested transcriptomic differences in three separate contexts. In the first, we compared MDA-MB-231 cells treated with doxorubicin to control MDA-MB-231. We found that these populations had easily distinguished morphologies, and our model was able to achieve a high classification accuracy. In the second context, we compared two separate subpopulations within the MDA-MB-231 cell line, and in the third, we compared two separate subpopulations within the MDA-MB-436 cell line. In both cases, the model was able to identify subpopulation identity with at minimum modest results.

Additionally, we interrogated the effect of cell orientation and interaction by increasing the area around each that cell that was input into the CNN. We found that increasing the bounding box initially improves the CNN’s classification ability, reaching an optimal size followed by a drop off in discrimination ability. To further understand this phenomenon, we performed several *in-silico* experiments which either completely removed the surrounding around each cell or removed textural information about the cell. Even without this information, the CNN was able to correctly classify each subpopulation with moderate discrimination ability in each case. In both experiments, the contour of the cell of interest is emphasized by a sharp contrast between background and cell area. This supports recent research which has utilized active shape models for prediction^17,18,41^. However, we find that while both texture and the surrounding environment are important, including both enables the best discrimination ability.

One unexplored avenue for cell morphological analysis is the use of unsupervised approaches^42,43^ to monitor changes in cell populations. The ability to autonomously observe shifts and correlate them to specific events could enable researchers to observe cell health, view the effects of different drug mechanisms, and observe phenotypic plasticity without the need for techniques such as sequencing or fluorescent labeling. One method that has been used on cell populations is autoencoders^16,21,22,44^, which contain a reduced latent space that is representative of the input and have been used for unsupervised learning approaches. Autoencoders have been used primarily to perform latent space predictions and provide explanations for changes in cell morphology. Using this latent space to make inferences about changes in the transcriptome without *a priori* knowledge about this change could give insight into these changes without the need for sequencing.

While these studies were performed on cells that also included fluorescent labels, classifying phenotype using only phase contrast images opens multiple possibilities for longitudinal experiments. The first is that it can be extended to many populations. Many automated imaging platforms capture red and green fluorescence, such that a maximum of 4 population labels (red, green, yellow, and non-fluorescing cells) can be monitored simultaneously. Given sufficient differences in morphology, a CNN could be adapted to identify additional subpopulations without fluorescent tags. Furthermore, some populations might not be suited to stable integration of a reporter gene, but a deep learning algorithm does not require this because it learns from morphology and interaction. Finally, not using fluorescent markers to track subpopulations frees the operator to have fluorescent indictors available for other uses. For example, cells could be stably integrated using the FUCCI^45^ system, enabling the ability to monitor the growth of each subpopulation as well as cell cycle.

The use of morphology for the tracking of subpopulations provides an avenue for better understanding of the ecology and evolution of subpopulations. Recent research has demonstrated that mathematical modeling can be used to provide insights into more optimal treatment schedules^46^. However, these models are limited in their ability to consider the heterogeneity that exists within a tumor population. Using cell morphology for cell state identification can provide insight into the transcriptomic composition of a population when testing and identifying new therapeutic strategies and targets.

## Methods

### Cell Culture

MDA-MB-436 cells were cultured in high glucose DMEM (Sigma D5796, 4.5 g/L glucose), 10% FBS, 0.01 mg/mL insulin (ThermoFisher, 12585014), 16 µg/mL glutathione (Sigma G013), penicillin-streptomycin (Thermo Scientific, 15140122). MDA-MB-231 were cultured in high glucose DMEM (Sigma, D5796) containing 10% FBS and 1X penicillin-streptomycin (Thermo Scientific, 15140122). Cells were maintained at 37 degrees C in a 5% CO_2_ atmosphere.

### Drug Treatment

Doxorubicin hydrochloride (Cayman Chemical 150007) was reconstituted in water. Cell culture media was replaced with new media containing 550 nM doxorubicin. After 48 hours, drug-free media replaced the doxorubicin dosed media.

### scRNA-seq Library Preparation

MDA-MB-231 samples from untreated and treated populations were harvested. Cells were loaded into wells of a Chromium A Chip. Libraries were prepared using the 10XGenomics 3’ single cell gene expression (v2) protocol. Paired end sequencing was performed using a NovaSeq6000 with an S4 chip (400 cycles) according to the manufacturer’s instructions.

### scRNA-seq Analysis

MDA-MB-231 single cell data was aligned to the GRCH38-2020-A and processed using Cell Ranger v7.1. All data was post-processed in the same way as described previously^30^ barring cell cycle regression. To make comparisons across samples and subsamples, we computed Pearson Correlation Coefficients (PCC) between each sample. To do so, we integrated all samples by re-normalizing data after concatenation. We then calculated the top 50 principal component values and found the PCC of these principal component values between each sample.

### Fluorescent nuclear labeling of MDA-MB-231 Cells

MDA-MB-231 cells were stably labeled using a sleeping beauty transposase system. MDA-MB-231 cells were grown to 90% confluence in a 10cm dish, then transfected with Lipofectamine 3000 according to the manufacturer’s protocol with 2.5 µg of a plasmid containing Sleeping Beauty transposase (SB100X, AddGene #34879) and 2.5 µg of a plasmid containing either GFP or mCherry on a nuclear localization signal (NLS) and flanked by inverted repeats. Stably transfected cells were selected for after 72 hours with G418, then expanded for two passages before antibody staining.

### Antibody staining and cell sorting

GFP and mCherry labeled MDA-MB-231 cells were stained with 2 µL ESAM-PE-Vio770 (Miltenyi, 130-115-039) per million cells. Cells were gated on cell single cell fraction, fluorescent channel, and ESAM staining. ESAM-low cells were sorted from the mCherry labeled population and ESAM-high cells were sorted from GFP labeled population. Enriched populations were expanded for at least two passages then further purified by performing another round of staining and FACS.

MDA-MB-436 cells were resuspended in cell staining buffer (PBS + 5 mM EDTA + 1% BSA + 1.6 mM NaOH + 0.01% sodium azide) and incubated in 1:100 diluted Zombie Violet viability dye (Biolegend, 423113) for 5 minutes on ice, then incubated with the 9 µL of APC-conjugated CD317 (BST2) antibody (Miltenyi, 130-101-660) per 1e6 cells for 20 minutes. Stained cells were washed 3 times with cell sorting buffer supplemented with 1:1000 diluted Zombie viability dye, then passed through a 40 µm cell strainer before FACS. Collection media for live cell sorting was prepared by supplementing complete media with 25 mM HEPES. Cells were sorted into 15 mL tubes containing 7 mL of collection media with gating on live cells, FSC-A/SSC-A, single cells, and BST2-lo or BST2-hi expression. Collected cells were spun down at 300 x g for 10 minutes, then plated in 50% conditioned media for 24 hours before transitioning to fresh, complete media. Conditioned media (CM) was prepared from complete media incubated on a 70% confluent plate of parental cells for 24 hours. CM was spun down at 500 x g for 10 minutes and supernatant was passed through a 0.22 µm filter. CM was diluted to 50% with fresh, complete media. Sorted subpopulations were allowed to recover from the sort and expand for a week before analysis.

### Image Acquisition and Processing

All images were acquired on an Incucyte S3. All cells were acquired at 20X magnification. For 20X magnification images, 9 images were taken per well. Fluorescent composite images were only used for cell phenotype ground-truth identification. To reduce the total number of cells per image, each 20X image was split into 16 separate image patches.

### Cell Segmentation

Image segmentation for cell lines in low to medium confluency images was achieved by training a custom Mask R-CNN instance segmentation algorithm. To obtain ground-truth segmentations, we first used Cellpose, a generalist segmentation algorithm, to obtain initial masks. We then reviewed each segmentation generated by Cellpose and augmented them when necessary to achieve better segmentation. Training by Cellpose alone was not effective, so these masks were fed into Mask R-CNN for specialized training. MDA-MB-231 cells had an associated precision (AP) score of 66%. MDA-MB-436 cells had an AP score of 55%.

### Cell Phenotype Discrimination Using a CNN

For each experiment, we chose to use a pre-trained network with residual connections (ResNet-152^25^) as they had shown promise in image classification. We used a stochastic gradient descent optimizer with an initial learning rate of 0.05 that decayed on each epoch. Any predefined augmentations (such as blacking out the background, increasing the surrounding bounding box, etc.) were applied to each image, and then were centered within a 300×300 pixel black background for 20X magnification images. To avoid overfitting, images were randomly flipped and rotated each epoch. Each image was normalized to the dataset mean and standard deviation. Models were trained for 30 epochs or until accuracy on the test set did not increase for 5 epochs. A hold-out well was used as a test set for subpopulation identification, and one well from each plate in the untreated and treated MDA-MB-231 experiments was used as a test set for those comparisons.

## Data availability

Images and models used in this paper are available upon request.

## Code availability

Scripts for training, testing, and segmentation are available at www.github.com/brocklab/transcriptomicCellMorphology

## Acknowledgments

We thank R01CA226258, R01CA255536, and U01CA253540 (to AB) which provided funding that supported this project. 10X Genomics Single Cell 3’ Gene Expression and TagSeq preparation were performed by the Genomic Sequencing and Analysis Facility at UT Austin, Center for Biomedical Research Support (RRID: SCR_021713). Flow cytometry and FACS was performed at the Center for Biomedical Research Support Microscopy and Imaging Facility at UT Austin (RRID: SCR_021756). TJ thanks Clarence Yapp for guidance on instance segmentation, model training, and image augmentation.

