## Supplementary Information for "Deep learning identifies heterogeneous subpopulations in breast cancer cell lines"

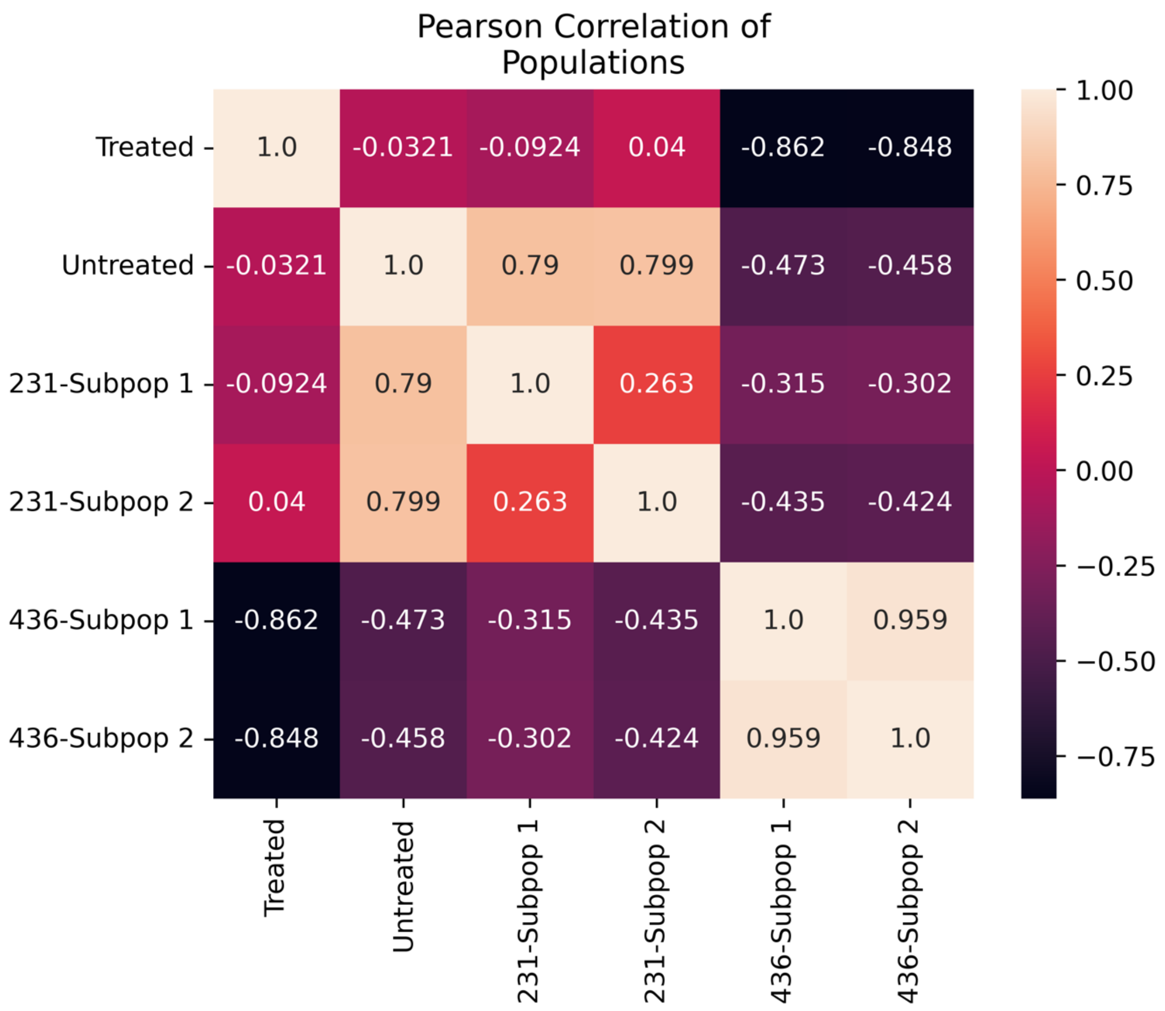


**Supplementary Figure 1.** Pearson correlation coefficient values calculated using 50 principal components from concatenated transcriptomic data.
